# Two distinct green algal symbionts cohabiting in the Japanese black salamander *Hynobius nigrescens*

**DOI:** 10.64898/2026.01.07.698294

**Authors:** Baptiste Genot, Swapnil Eishraq Abedin, Naoshi Shinozaki, Ryan Kerney, Shinichiro Maruyama

**Affiliations:** Department of Integrated Biosciences, Graduate School of Frontier Sciences, The University of Tokyo, Kashiwa, Chiba 277-8562, Japan; Japan Amphibian Laboratory, Nikko, Tochigi, Japan; Gettysburg College, Gettysburg, PA, U.S.A.

## Abstract

Green algae in the genus *Oophila* are associated with eggs of amphibians, e.g. salamanders and frogs, and are the only known photosynthetic endosymbionts in vertebrates. However, phylogenetic placements of “*Oophila*” algae associated with amphibians into two distinct clades have caused significant confusion and hindered our understanding of the nature and diversity of this symbiotic system. To gain insights into this unexplored symbiotic model, we carried out a sampling of the Japanese black salamander *Hynobius nigrescens* eggs, an endemic species in Japan, covering five distinct locations. Egg masses from *H. nigrescens* all contained green algae, some of which were sampled, isolated, and cultured. In total, 32 algal isolates could grow as free-living cultures and were used for phylogenetic and morphological analyses. Our results supported the co-existence of two *Oophila* subgroups with *H. nigrescens* embryos. One is associated with the known subclades of *O. amblystomatis* found in North America, and the other belongs to an *Oophila/Chlorococcum* group in the Stephanosphaerinia. The isolates in each subgroup showed identical morphology across sampling sites, consistent with our phylogenetic findings. These new, unique cultures of amphibian-associated green algae will help to understand photosymbiosis between vertebrates and algae.

## Introduction

Photosymbiosis is a natural interaction between a non-photosynthetic host and a photosynthetic symbiont. Perhaps the most iconic example of these remarkable interactions is the symbiosis between dinoflagellate algae and animals belonging to the phylum *Cnidaria* [1,2], including the reef-building corals. In freshwater environments, green algae species from the genus *Chlorella* are known to be symbionts of some host species (the cnidarian *Hydra viridissima*, the amoeba *Mayorella viridis*, the heliozoan *Acanthocystis turfacea*, and the ciliate *Paramecium bursaria* [2– 4]). All these examples include invertebrate hosts, and the photosymbiosis is extremely rare in vertebrates that developed adaptive immunity, involving specialized cells. As an exception, green algae belonging to the *Oophila* species in North America are known to be amphibian “egg-loving” photosymbionts. They are primarily associated with the yellow spotted salamander *Ambystoma maculatum*, and naturally invade egg masses during early embryo development. Known for more than 130 years [5], *Ambystoma maculatum* and *Oophila amblystomatis* interactions have been studied as the sole model of photosymbiosis in vertebrates. Algae proliferating in egg capsules were also found in embryonic tissues and incorporated into animal cells [6,7]. This led to two models of “ectosymbiosis” outside of the animal (in egg fluid) and “endosymbiosis” (or intracellular) inside embryos and adults of *Ambystoma maculatum*. In *A. maculatum* egg capsules, *O. amblystomatis* was shown to be beneficial for embryo development, generating oxygen as a product of photosynthesis and thus limiting hypoxia in egg masses [8–11], while symbionts would benefit from host nitrogen and other waste, suggesting a mutualistic relation. Meanwhile, intracellular interactions between algae and animal cells have been highlighted through transcriptomics [6]. Notably, the algae shifted toward fermentation, while the host cells exhibited markers of immune suppression, suggesting a parasitic, rather than mutualistic, aspect of the relationship. *Oophila* algae were also found in other eggs of salamander and frog species (reviewed in [12], *Ambystoma gracile* (Northwestern Salamander), *Rana sylvaticus* (wood frog), and *Rana aurora* from North America [13]. In Europe, two species of frog, *Rana dalmatina and R. temporaria*, showed algal presence in eggs [14,15]. Lastly, *Oophila* algae were reported in Japan, associated with *Hynobius nigrescens* (Japanese black salamander) [16]. However, no endosymbiotic algae have been reported on these species. Phylogenetic analyses of *Oophila* spp. have been demonstrated in several studies, but resolving green algae, especially chlamydomonadales phylogeny, is complex and would require more sampling efforts to understand the organization and the evolution of this genus. First, *Oophila* spp., as green algae associated with amphibian eggs, can be classified into two distinct polyphyletic clades (Clades A and B). As contradicting phylogenetic studies were issued [15,17–19], the discovery of *Oophila* species in different hosts and different continents could help to clarify the complexity of the *Oophila* phylogeny. *Oophila/Chlorococcum* “ Clade A” regroups the algae phylogenetically clustered with *Chlorococcum* free-living species but is found in amphibian eggs (including samples from *A. maculatum* and *Rana dalmatina*). *Oophila amblystomatis* “Clade B” regroups the endosymbiotic and ectosymbiotic symbionts from North America. It also includes egg symbionts found in *A. gracile* and the frogs *R. auror*a and *R. sylvaticus*, which are not found inside animals in these species. As presented in [13], *Chlamydomonas moewusii* is one of the closest free-living relatives and serves as a reference point for distinguishing symbionts belonging to clade B.

Looking at the Japanese *Oophila* identified in 2017, the five partial sequences obtained formed a Japanese subclade J1, nested in the *Oophila* clade B [16]. Divergence between the yellow spotted salamander and the Japanese black salamander occurred about 200 MYA [20], indicating that algal–amphibian associations in Japan and North America have evolved independently for a sufficiently long period to allow distinct evolutionary trajectories. As of today, no deposited J1 clade *Oophila* cultures exist, and five partial 18S DNA sequences have been used to map the J1 clade position in chlamydomonadales [16]. In this study, we conducted a sampling campaign in Japan, aiming to collect, isolate, and further examine the Japanese *Oophila* phylogenetic position among green algae. 32 isolates were cultured and mapped in the current phylogenetic tree, showing a Japanese-specific sub-clade A (related to the genus *Chlorococcum*) and sub-clade B (nested between the known *Oophila amblystomatis* Clade B subclades III and IV) of amphibian-associated algae. Our data support that *H. nigrescens* eggs are inhabited by two distinct green algae lineages belonging to the polyphyletic group of green algae regrouping *Oophila amblystomatis* and *Chlorococcum* algae.

## Results and Discussion

### Algae associated with H. nigrescens egg masses are likely green algae belonging to the genus Oophila

Clutches were collected from five distinct locations in Ishikawa, Toyama, and Tochigi areas, in Japan (Figure 1A). Single-cell algae were isolated from egg fluids and egg capsules (see material and methods), leading to 32 culture isolates (four to eight isolates per site from 5-6 eggs). During sampling, egg and algal morphology was assayed using a fluorescence stereomicroscope and an optical microscope. All clutches and eggs were positive for algal presence, and all clutches were open at an embryo development stage estimated between 35 and 37 [21]. *H. nigrescens* masses were all cloudy (as seen in Figure 1C). *H. nigrescens* eggs can be found white, clear, or intermediate and environmental conditions such as altitude or maximum show depth were shown to be important factors explaining this variation [22]. All eggs showed a green coloration under white light (Figure 1C) and a typical red chlorophyll fluorescence when observed under fluorescence (Ex/Em 540-580/610 nm) (Figure 1A). Non-motile algae embedded in the egg capsule and motile algae in the egg fluids were observed (Figure 1D, Supplemental file 1). Optical microscopy confirmed the presence of cells with a crown-shaped cell wall (Figure 1D), consistent with previous observations and characteristic of J1 *Oophila* (Clade B, Japanese subclade) (described as “reticulate protuberance” in [16]. Free-swimming flagellated cells were also observed, but couldn’t be identified further using optical microscopy.

**Figure 1.**
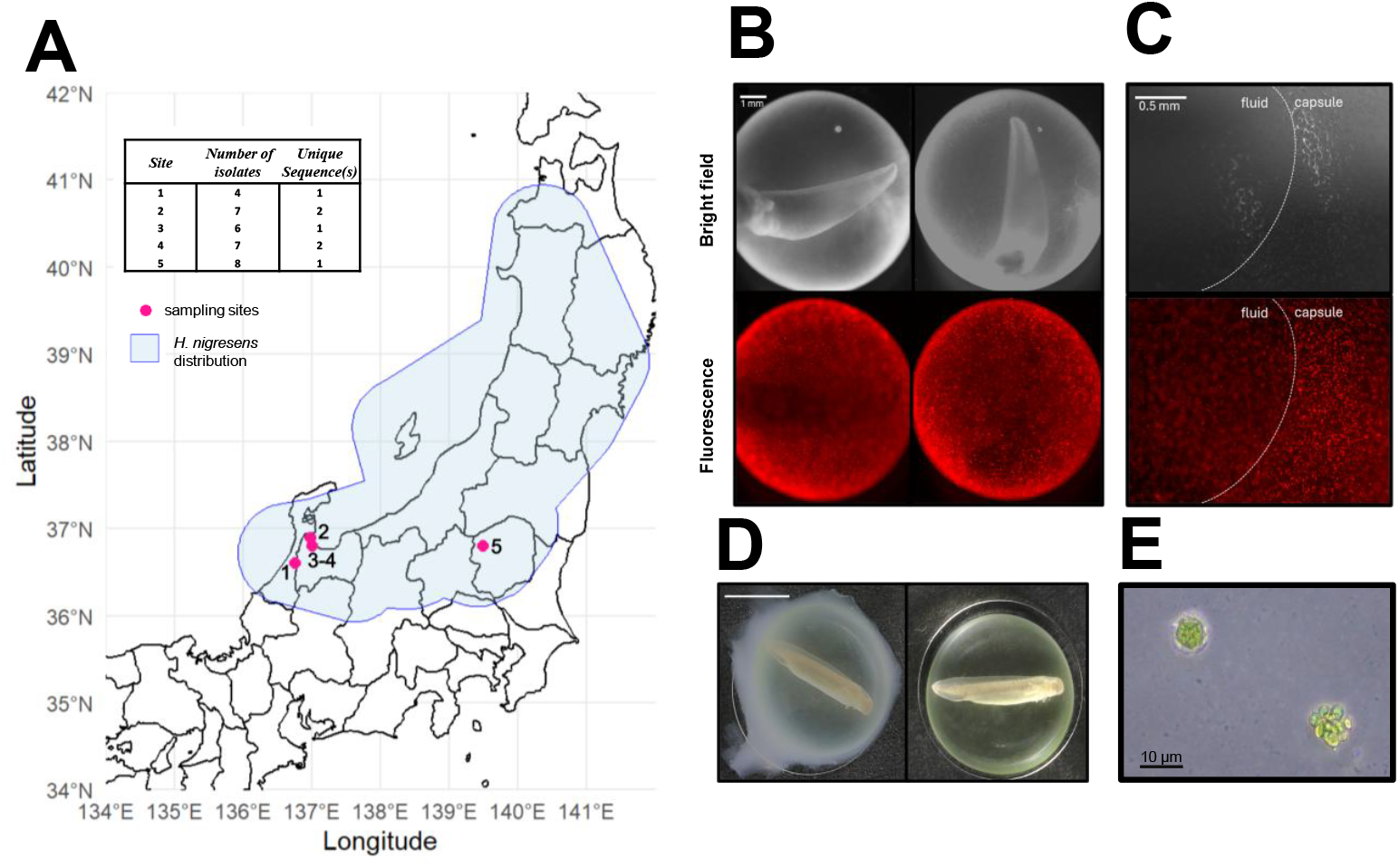
A. Sample information of *Hynobius nigrescens* egg masses. Observation records of *Hynobius nigrescens* in iNaturalist records available as of September 2025 were mapped as the putative distribution. B. Bright-field (top) and fluorescence micrographs (bottom) of *H. nigrescens* eggs (scalebar = 1 mm). C. Zoomed view of *H. nigrescens* eggs (scalebar = 0.5 mm). Egg fluid and egg capsule are indicated and separated by a dashed line. Chlorophyll was observed at Ex/Em 540-580/610 nm. D. Magnified views of the border between the fluid and the egg capsule (scalebar = 0.5 mm). E. Representative micrographs of the sample preparation before and after cleaning. E. Green algae sampled from an egg capsule (scale bar = 10 μm).

### Phylogenetic analysis helped to clarify the Oophila subclade positions among green algae

To attribute a phylogenetic position to the 32 isolates, 18S DNA was sequenced using universal eukaryotic primers [23]. The sequence alignment of the 32 isolates resulted in two unique sequences (Supplemental Figure 2). One sequence (28 occurrences) at all sites (Ishikawa, Kanazawa, and Nikko prefectures), and the other one (4 occurrences) at sites 1 and 4 (Ishikawa and Kanazawa prefectures). The two sequenced 18S regions were different (91.82% identity), implying that two potential *Oophila* strains exist in our samples. Phylogenetic analysis using known *Oophila* sequences, from all identified hosts and free-living isolated algae, was carried out using IQ-TREE (Figure 2) [24]. The unique sequence with 4 occurrences nested inside Clade B, and could be associated with the subclades III and IV described in [13]. These subclades contain *Oophila* algae found in *A. maculatum, L. aurora*, and *L. sylvaticus*. Our second 18S occurrence (28 sequences) was nested within Clade A, including *Oophila/Chlorococcum* identified in amphibian eggs, and the free-living *Chlorococcum* algae, *Hamakko caudatus, Haematococcus zimbabwiensis, Stephanosphaera pluvialis*. These observations support that clade A symbionts exist in Japan and are associated with salamander eggs. Other known sequences published as *Oophila* “clade A” and found in *A. maculatum* eggs were not clustered with the ones from *H. nigrescens-*associated algae (98.59% sequence identity vs. KY091670 and KJ635658; 97.35% sequence identity vs. KJ635657). These suggest that Japanese clade A algae are genetically distinct from North American clade A algae associated with the Yellow-spotted salamander.

**Figure 2:**
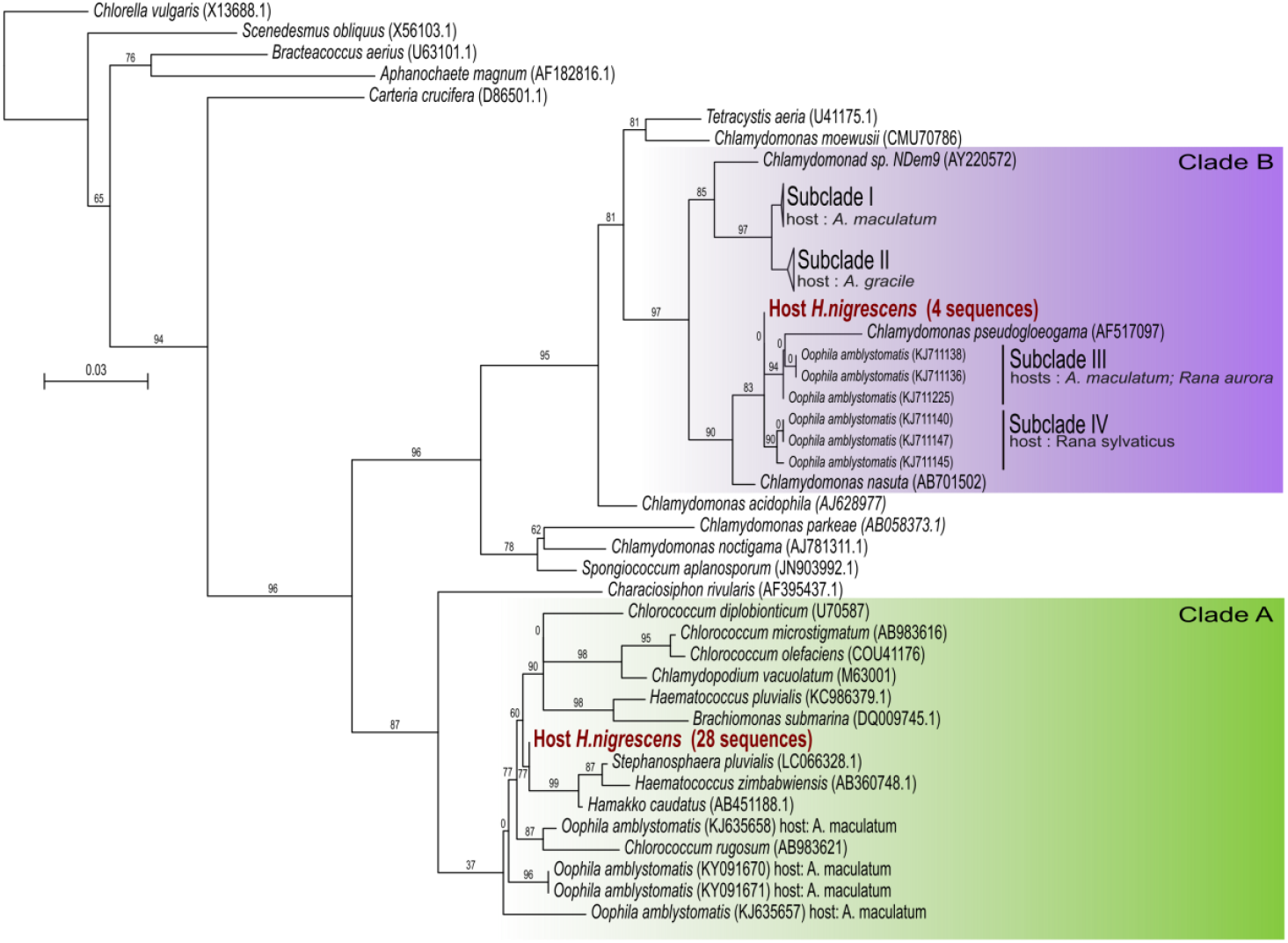
Maximum Likelihood phylogenetic tree of 18S rDNA sequences. Branch support values correspond to the standard nonparametric bootstrap (1000 replicates). SH-aLRT and aBayes posterior probability were also calculated (supplemental file 2). Collapsed *Oophila* subclade I and II include KJ711220, KJ711137, and KJ711190 accessions and KJ711161, KJ711173, and KJ711151 accessions, respectively. The animal host is indicated where the sequence is from an ectosymbiotic or endosymbiotic alga.

### Clade A and Clade B algae from H. nigrescens are morphologically different

After phylogenetic analysis, we successfully established five representative algal cultures of *Oophila* diversity in Japan. As phylogeny (Figure 2) showed two unique sequences, we deposited two cultures at NIES [to be added later]. These represent the first publicly available culture of *Oophila* from Japan. To compare the morphological characteristics of each algal isolate, we observed all five strains under a microscope and measured their cell sizes (Figure 3). The observations revealed that strains belonging to Clade B were larger than those belonging to Clade A. The median Feret diameter (also known as maximum caliper) of Clade A isolates was 7.23 µm at Site 4, 7.54 µm at Site 1, and 7.57 µm at Site 5, showing a consistent trend within the clade (Figure 3B). Similarly, the median Feret diameters of Clade B isolates were 10.8 µm at Site 4 and 11.7 µm at Site 1, also showing internal consistency but with larger cell sizes than those of Clade A. To test whether the median cell sizes differed significantly between Clade A and Clade B algae, we used a generalized linear model (GLM) to examine the effect of clade identity and sampling site on the Feret diameter of the cell. The results showed that both Clade (p < 2 × 10^−16^) and Site (p = 8.59 × 10^−5^) had significant effects on the Feret diameter. In addition, *Oophila/Chlorococcum* Clade A isolates were generally more spherical in shape (Figure 3A). Site 1 and Site 4 are located in Kanazawa, Ishikawa prefecture, and Toyama, respectively, and Site 5 is located in Nikko, Tochigi Prefecture. Despite these geographically distinct sampling sites, isolates within the same clade exhibited similar morphological features and phenotypes. This morphological consistency supports the genetic separation of Japanese *Oophila* into two different lineages. Furthermore, because all five isolates were directly obtained from *H. nigrescens* eggs, our results support that multiple *Oophila* lineages can coexist within the same host species.

**Figure 3:**
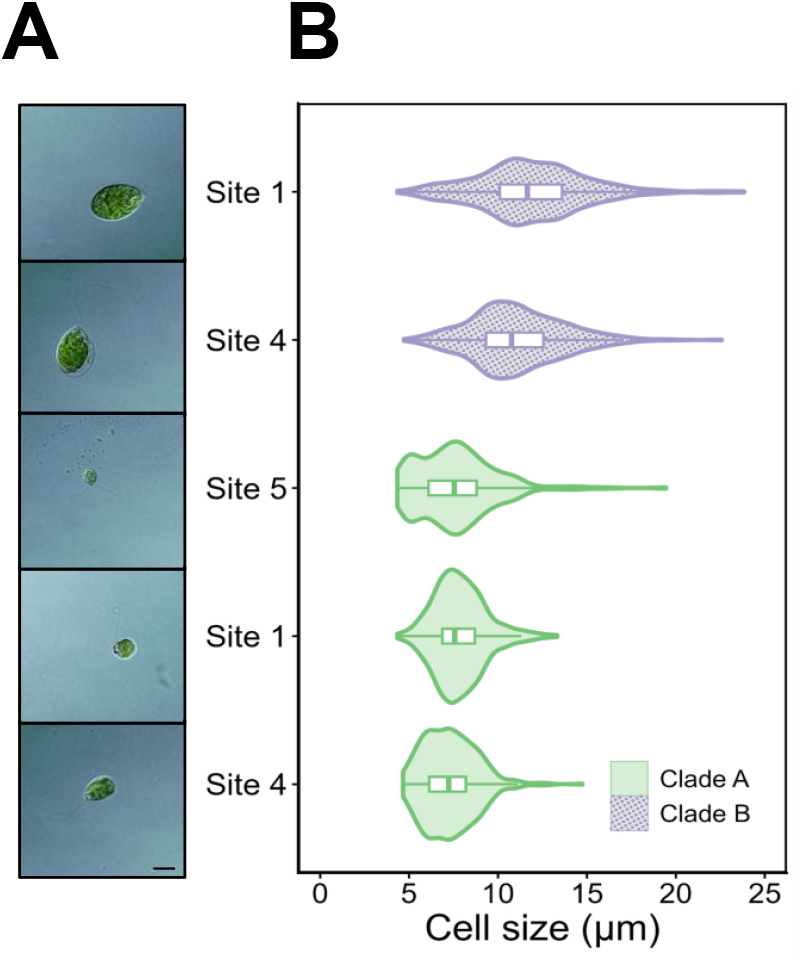
Representative micrographs of five strains of *H. nigrescens* egg algal symbionts. Each image corresponds to the violin plot. All photos were taken using differential interference contrast at 100x magnification. Scalebar = 10 μm. C. Cell size distribution of five *H. nigrescens-associated* algal strains. The X-axis represents the Feret diameter (maximum cell diameter) of isolates. For each isolate, 500 cells were measured. Statistical significance was assessed using a generalized linear model (GLM) with a Gaussian family, including Clade, Site, and their interaction (Feret diameter ∼ Clade × Site; Clade: p < 2 × 10^−16^; Site: p = 8.59 × 10^−5^).

### Deeper sampling allowed to unravel the complexity and diversity of amphibian-associated algae

As *H. nigrescens* is endemic to Japan, we expected to unravel the diversity of green algae associated with egg masses by sampling different locations in the known habitat of the black salamander. We isolated 32 potential *Oophila* green algae and used the 18S sequence to determine the phylogenetic positions of two strains that occurred 4 and 28 times, respectively. No other algal species were isolated from egg fluids or capsules during our experiments. The Japanese Clade A algae identified in this study were not clustered with other *Oophila* species, but instead with free-living relatives such as *Hamakko caudatus*. The latter was identified in Japan [25] and might indeed be the closest relative of amphibian-associated algae. Together with *Haematococcus zimbabwiensis* and *Stephanosphaera pluvialis*, all these algae form the Stephanosphaerinia group, which is distinctly nested among other green algae [26]. It is important to note that these groups are not yet fully defined, and our current understanding of their phylogeny will likely be revised as additional species are discovered and new transcriptome-based analyses become available [19]. Regarding the identification of Clade B algae, our results and the previous study [16] confirmed the existence of the subclade J1, the Japanese subclade *O. amblystomatis*. Unfortunately, we chose not to include published sequences as they were too short and would reduce our phylogeny quality. Clade B algae were previously identified, but no cultures were available. In this study, we successfully established Clade B cultures (along with Clade A algae) that can be used in future experiments and in taxonomy studies. Clade B isolates in *H. nigrescens* and *A. maculatum* are closely related but genetically distinct from each other. Deeper genomics and transcriptomics would help to understand how the *Oophila* species co-evolved with two salamander species. *Ambystoma maculatum*, the North American yellow-spotted salamander, is known to associate with *Oophila amblystomatis* in clade B, as reported by various studies [6,7,13]. *A. maculatum* and *O. amblystomatis* have a particular interaction, as the algae can invade egg fluids and embryo cells. While this unique model of algal-vertebrate symbiosis has been studied extensively, Clade A algae have also been less frequently detected in *A. maculatum* eggs [14,18] and no endosymbiosis has been reported. A recent study [27] even suggested that no chlorophytes outside the clade B can be found in egg fluids, using 18S reads from a vast array of samples. Our results showed that Clade A algae are found more frequently (using single-cell isolation and culture) in *H. nigrescens*. Clade B was also identified, suggesting that two different lineages of green algal symbionts cohabit in *H. nigrescens* eggs. These two strains were morphologically different, Clade A being smaller in diameter in culture compared to Clade B. At last, we did not observe algal presence in animal tissues, and so far, no intracellular algae have been reported in *H. nigrescens* embryos. The unexpected co-occurrence of both symbiont lineages reveals new biological insights and makes *H. nigrescens* a unique model of photosymbiosis. In this study, egg mass jelly was not sampled as it could contain free-living algae that are not associated with eggs or egg fluids. It is now interesting to look at how and when these two symbionts can sense and enter masses to reach the eggs. The ecological basis of this symbiosis also merits investigation. The fact that two algal species coexist within salamander eggs indicates that they do not negatively affect one another and that the host likely offers a shelter, potentially protecting them from predators, pathogens, or abiotic stresses. To get more insight into algal diversity in *H. nigrescens* eggs, techniques such as DNA metabarcoding [14] or transcriptome analysis [27] should be carried out in the future. The Clade A/B ratio in Japanese salamander eggs seems to be different from that in the North American yellow-spotted salamander. As the two species diverged 200 MYA, interesting questions emerged regarding the evolution of the two systems: why is Clade A dominant in Japan but nearly absent in North America, and conversely for Clade B? What about other amphibian eggs (e.g., *R. japonica* shares a similar distribution with *Hynobius nigrescens* and is also endemic to Japan)? Additional sampling and taxonomy efforts, in combination with molecular and cellular biological experiments, will help to understand the ecological importance of the genus *Oophila*.

## Material and methods

### Sampling Hynobius nigrescens clutches in Toyama/Kanazawa prefectures

Clutches (Sites 1, 2, 3, and 4; Figure 1, Supplemental Figure 1) were found in several agricultural reservoirs on March 20^th,^ 2025, and washed and sampled in laboratory facilities on March 21^st,^ 2025. Sites coordinates were as follow: Site 1 (36.6108633,136.7480953 ; Ishikawa prefecture); Site 2 (36.8995992,136.9801988; Toyama prefecture); Site 3 (36.8141971,137.0075838; Toyama prefecture); Site 4 (36.8074779,137.0101021; Toyama prefecture)

### Sampling Hynobius nigrescens clutches in Nikko prefecture

Clutches (Site 5; Figure 1, Supplemental Figure 1) were found in an artificial breeding pond on June 5^th,^ 2025, and were washed and sampled the same day in laboratory facilities. Site 5 coordinates were:36.80178,139.49053.

### Sample map

Mapping of the sampling campaign was realized using iNaturalist [28] resources and the code available in Supplemental file 5. Packages used are referenced as[29–33].

### Sample handling and preparation

Clutches were washed three times with Milli-Q water and 70% ethanol using a strainer. All leaves and other organic materials were carefully removed before transferring them into a plastic container in 20% Holtfreter’s solution. 100% Holtfreter’s stock solution was made using 3.46 g/L NaCl, 0.05 g/L KCl, 0.1 g/L CaCl2, 0.2 g/L NaHCO3, and pH adjusted to 7.4. Each clutch was handled separately, and the sampling area was thoroughly washed after each completed algal sample. Using a sterile plastic pipette and a razor blade, clutches were opened, and a few eggs were separated and cleaned from the mass jelly. Cleaned eggs were then washed three times in 20% Holtfreter’s solution baths. Using a syringe, egg fluid was collected and placed in a 10 mL AF6 [34] culture flask. The remaining egg capsule was washed three times in 20% Holtfreter’s solution baths and placed in a 10 mL AF6 culture flask. On the following day, single-cell algae were isolated by pipetting into 24-well plates containing 500 μL AF6 and incubated for 15 days at 15°C in a 12-hour daylight cycle incubator. If growth was observed, cells were then transferred to 30 mL culture flasks.

### Molecular biology and phylogenetics analysis

To extract DNA, 500 μL of cells were collected by centrifugation (10,000g, 5 minutes) and used as input for the Maxwell RSC with the plant DNA kit protocol. 18S/ITS2 region was amplified by PCR using the forward primer SR1(5’-TACCTGGTTGATCCTGCCAG-3’) and the reverse primer SR12 (5’-CCTTCCGCAGGTTCACCTAC-3’)[23]. The DNA polymerase Mighty ampV3 was used following the manufacturer’s protocol and using the following program: 98°C 2 min ; [98°C 30sec, 55°C 30sec, 68°C 3min]x30; 68°C 3min. PCR products were then purified using a Wizard SV Gel and PCR Clean-Up System and sent for Sanger sequencing (Eurofins) using the SR12 primer.

### Phylogenetic tree

18S sequences were aligned using MAFFT [35] and trimmed (gap threshold 0.8) using trimAl [36] before being manually verified (suppl. File 2). IQ-TREE version 3 was then launched in AUTO mode with the following parameters: Non-parametric bootstrap with 1000 replicates; Approximate Likelihood-Ratio Test (aLRT) with 1000 replicates; Approximate Bayesian support. The best-fit model output was HKY+R3 (Hasegawa–Kishino–Yano model + FreeRate model with 3 rate categories). TreeViewer [37] was used to shape the final tree. Accessions used in the figure are as follows: *Chlorella vulgaris* (X13688.1), *Scenedesmus obliquus* (X56103.1), *Bracteacoccus aerius* (U63101.1), *Aphanochaete magnum* (AF182816.1), *Carteria crucifera* (D86501.1), *Chlamydomonas_acidophila* (AJ628977), Tetracystis aeria (U41175.1), *Chlamydomonas moewusii* (CMU70786), *Chlamydomonad sp. NDem9* (AY220572), *Chlamydomonas pseudogloeogama* (AF517097), *Chlamydomonas nasuta* (AB701502), *Oophila amblystomatidis* subclade IV (KJ711138, KJ711136, KJ711125), *Oophila amblystomatidis* subclade IV(KJ711140, KJ711147, KJ711145), *Oophila amblystomatidis* subclade I (KJ711220, KJ711137, KJ711190), *Oophila amblystomatidis* subclade II (KJ711161, KJ711173, KJ711151), *Oophila amblystomatidis* subclade III (KJ711138, KJ711136, KJ711225), *Oophila amblystomatidis* subclade IV (KJ711140, KJ711147, KJ711145), *Chlamydomonas parkeae* (AB058373.1), *Chlamydomonas noctigama* (AJ781311.1), *Spongiococcum aplanosporum* (JN903992.1), *Characiosiphon rivularis* (AF395437.1), *Chlorococcum diplobionticum* (U70587), *Chlorococcum diplobionticum* (U70587), *Chlorococcum olefaciens* (COU41176), *Chlamydopodium vacuolatum* (M63001), *Stephanosphaera pluvialis* (LC066328.1), *Haematococcus pluvialis* (KC986379.1), *Brachiomonas submarina* (DQ009745.1), *Haematococcus zimbabwiensis* (AB360748.1), *Hamakko caudatus* (AB451188.1), *Chlorococcum rugosum* (AB983621), *Oophila amblystomatidis* (KJ635658, KY091670, KY091671, KJ635657). Sequences alignment and IQ-TREE.log file are included in the supplemental file 2.

### Cell pictures and size

*Oophila* cells were fixed in 4% paraformaldehyde (PFA) diluted in phosphate buffer (PBS). and incubated at 4°C for 10 minutes. Samples were then centrifuged (3000 g, 5 minutes) and most of the supernatant removed. Algae were then transferred into a glass slide and covered with a coverslip (for DIC observation) or to a glass-bottom dish (for imaging with the CSU system) prior to observation. DIC images were acquired using an Olympus BX53 upright microscope. Observations were performed under bright-field conditions using a 100× oil objective lens with a DIC prism. The exposure time was set to 100 ms. For cell counting, images were acquired using an Olympus CSU system (Yokogawa Electric) mounted on an Olympus IX81 inverted research microscope. Cells were photographed with a 10× objective lens and an exposure time of 100 ms. Cell counting and sizing were performed using Fiji [38]. The acquired images were converted to 8-bit format and subjected to averaging with a Radius of 1 for noise reduction. Particles were analyzed using the Auto Threshold function. Cell counts and the Feret diameter (maximum cell diameter) were measured for particles meeting the following criteria: size: 10−300 μm^2^, circularity: 0.7−1.0. For each strain, 500 cells were counted and measured from a single field of view. All statistical analyses and data visualizations were performed using R statistical software [39]. The effect of clade and sampling site on cell size (Feret diameter) was determined using a General Linear Model (GLM) assuming a Gaussian family, defined as: Feret diameter ∼ Clade*Site (including interaction term between clade and site). Violin and box plots were generated using the ggplot2[33] and ggpattern [40] packages.

## Data accessibility

Sequences analyzed in this study are available at the NCBI accessions PX558014 to PX558046, along with sample descriptions. Representative cultures of *Oophila amblystomatis* Clade B and *Oophila/Chlorococcum* Clade A were submitted to the National Institute for Environmental Studies (NIES) and will be available under references NIES XXXX and NIES XXXX.

## Acknowledgements

We thank Shintaro Seki for their help in sampling *Hynobius nigrescens* eggs. The computation was performed using the Research Center for Computational Science, Okazaki, Japan (Project: NIBB, 25-IMS-C233).

## Supplemental Figures

**Supplementary Figure 1.**
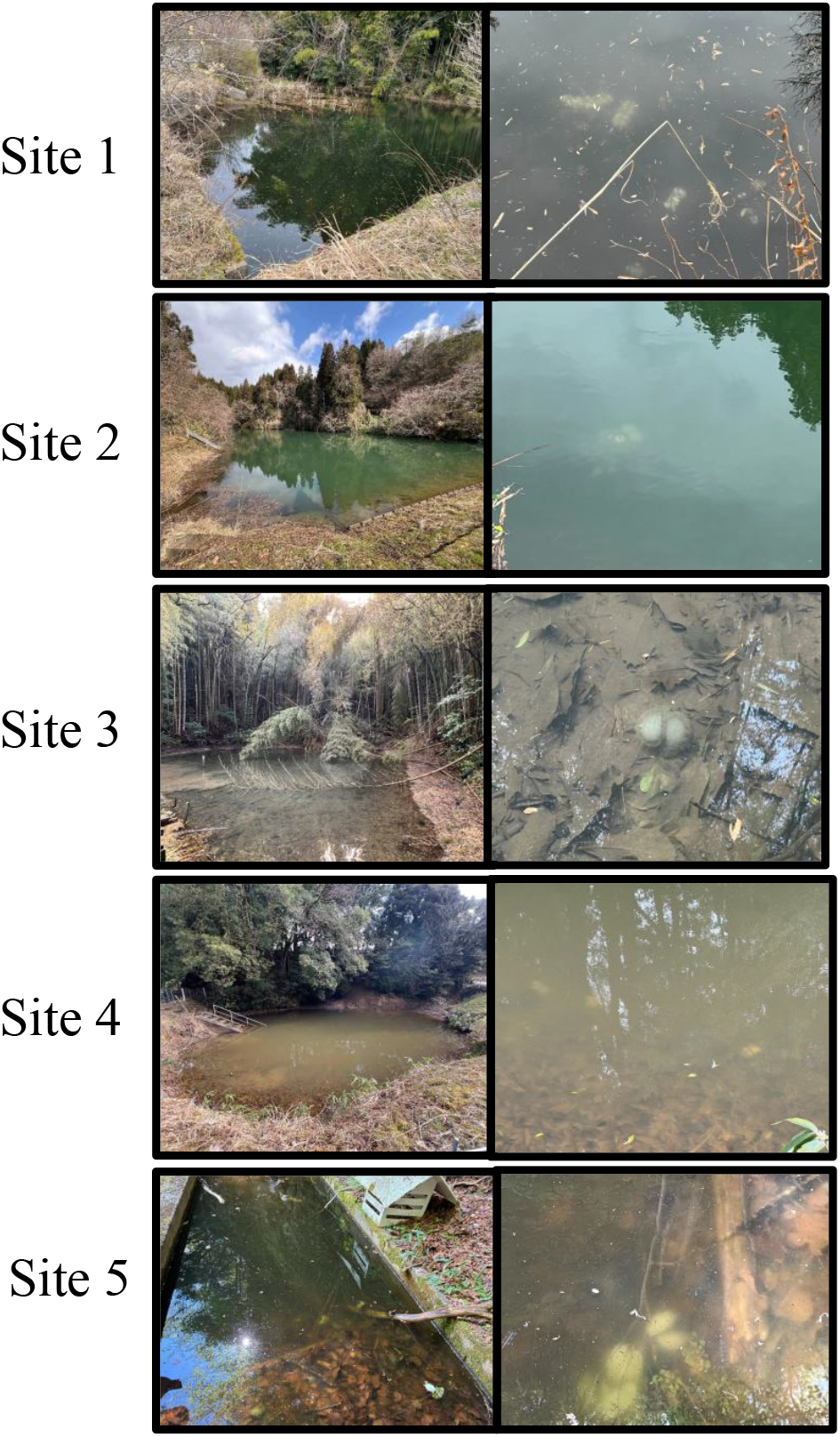
Photograph of the sampling sites (Authors of photograph: Shintaro Seki, Daisuke Yamagushi) and identification of green *Hynobius nigrescens* eggs.

**Supplementary Figure 2.**
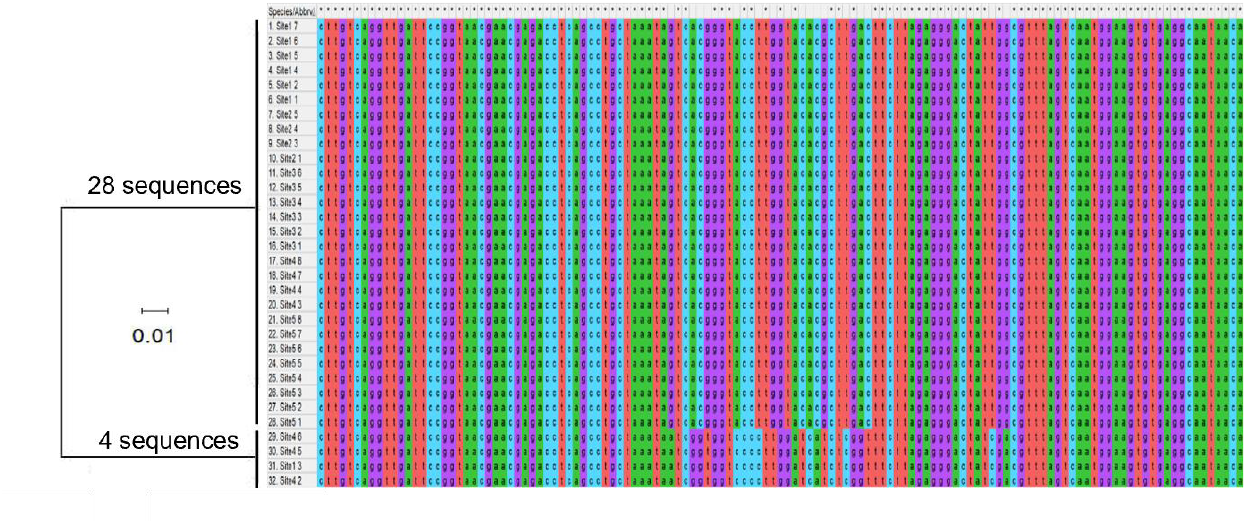
Neighbor joining tree and partial 18S rDNA sequence alignment of the 32 amphibian-associated algae isolates. MEGA12 [41] was used for creating the alignment and the neighbor-joining tree.

## Supplemental Files

Supplemental File1

Video of live green algae swimming inside *Hynobius nigrescens* eggs.

Supplemental File 2

Iqtree log and alignment files

Supplemental File 3

Unique sequences alignement

Supplemental File 4

R Code used for this study

